# Integrative Cross-Cohort Meta-Analysis Reveals a Conserved Dysbiotic Signature of *Streptococcaceae* and *Lachnospiraceae* in Multiple Sclerosis

**DOI:** 10.64898/2026.07.25.740724

**Authors:** Amaan Arif, Prekshi Garg, Prachi Srivastava

## Abstract

**Background:** Multiple Sclerosis (MS) is a chronic autoimmune disorder characterized by inflammation and demyelination in central nervous system (CNS). Although increasing evidence suggests that gut microbial dysbiosis contributes to MS pathogenesis through the microbiota-gut-brain axis, reproducible microbial signatures associated with disease progression across independent clinical cohorts remain incompletely characterized.

**Objective:** This study aimed to identify conserved gut microbial alterations associated with Multiple Sclerosis by integrating publicly available human gut microbiome datasets and characterizing disease-associated microbial signatures linked to immune dysregulation.

**Design:** Human gut metagenomic 16S rRNA sequencing data from MS patients and healthy controls obtained from publicly available repositories (NCBI, Bioproject). Raw sequencing reads were processed using a standardized microbiome analysis workflow, including quality control, denoising, taxonomic assignment, phylogenetic reconstruction, diversity analyses, and differential abundance testing. Microbial community structure was evaluated using alpha- and beta-diversity analyses, while statistically significant differences between study groups were assessed using PERMANOVA, Kruskal-Wallis, and ANCOM to identify disease-associated bacterial taxa.

**Results:** Integration of independent cohorts revealed consistent alterations in the gut microbial composition of MS patients compared with healthy controls. Significant reductions in microbial diversity and distinct microbial community structures were observed in MS. Differential abundance analysis demonstrated enrichment of the pro-inflammatory family *Streptococcaceae*, whereas beneficial short-chain fatty acid-producing taxa, particularly *Lachnospiraceae*, were significantly depleted in MS patients. These conserved microbial alterations indicate disruption of immune-regulatory bacterial communities and support the involvement of gut microbial dysbiosis in MS-associated neuroinflammation.

**Conclusion:** This study identifies a reproducible gut microbial dysbiosis signature associated with Multiple Sclerosis, characterized by expansion of pro-inflammatory bacterial taxa and depletion of beneficial SCFA-producing microorganisms. These findings strengthen the evidence supporting the microbiota-gut-brain axis in MS pathogenesis and highlight microbial community signatures that may contribute to future biomarker development and microbiome-based therapeutic strategies.

## Introduction

Multiple sclerosis (MS) is a chronic autoimmune disease, frequency of which reaches an epidemiological level with devastating neurological health outcomes [1]. MS predominantly affects the myelin protective covering of nerve fibres in brain and spinal cord with inflammation, demyelination within the Central Nervous System (CNS) [2,3]. Partial degradation of this insulating sheath impairs rapid conduction of electrical impulses along nerve cells/ fibres that manifests as fatigue, motor weakness (clumsiness), sensory disturbances (e.g. numbness and tingling in hands or feet), vision defects due to optic neuritis, coordination difficulties referred to ataxia, cognitive dysfunction [4]. Fatigue is common and disabling symptoms experienced by more than half of patients and is a major factor responsible for impairing quality of life [5]. Mobility may also be affected by weakened muscles and legs that are spastic and uncoordinated leading to difficulties walking or an increased risk of falling [6]. The presence of cognitive impairment and emotional disturbances are also often complicating the adjustment to this disease [7,8].

MS is most commonly present in women and men aged 20 to 40 and is more prevalent in women than men [9,10]. MS is a polygenic or multifactorial disease caused by genetic predisposition as well as exposure to environmental factors that induce an auto-aggressive immune response. Here, typical immune cells like T and B lymphocytes cross the blood-brain barrier (BBB) is a selective protective envelope that usually prevents chemicals from passing through to reach the CNS. In the setting of MS, immune cells mistakenly identify the myelin sheath (a product made by one’s own body) as an outside invader and this serves to kick off inflammatory processes that result in demyelination & axonal loss [11].

The gut microbiota is a diverse community of microorganisms that plays an important role in maintaining immune homeostasis and intestinal barrier integrity. Changes in gut microbial composition caused by factors such as diet, lifestyle, or medication can disturb this balance, leading to altered immune responses and increased inflammation. Growing evidence suggests that these changes may contribute to the development and progression of several immune-mediated diseases, including Multiple Sclerosis (MS). The gut-brain axis has gained considerable attention in recent years as a bidirectional communication network connecting the gastrointestinal tract and the central nervous system [12]. Through immune regulation, microbial metabolites, and maintenance of the intestinal barrier, the gut microbiota influences brain function and neuroinflammatory processes [13]. Disruption of this communication may promote abnormal immune activation and contribute to the pathogenesis of MS, making the gut microbiota an important area of investigation for understanding disease mechanisms.

Recent studies have shown that changes in the gut microbiota may influence both the development and progression of Multiple Sclerosis (MS). Patients with MS often show reduced bacterial diversity, changes in specific bacterial families, and altered immune responses [14]. These microbial changes may contribute to immune dysregulation by reducing beneficial bacteria that promote regulatory T cells (Tregs), which are important for maintaining immune tolerance and preventing autoimmune responses [15]. In addition, lower levels of short-chain fatty acid (SCFA)-producing bacteria have been reported in MS patients, suggesting a possible link between gut microbial imbalance and disease progression [16–18]. However, despite these findings, the gut microbial changes reported in MS remain inconsistent across different studies. Differences in study populations, geographic regions, sequencing methods, and data analysis approaches have resulted in variable microbial signatures. Therefore, it is still unclear which microbial changes are consistently associated with MS across independent patient cohorts. Identifying these reproducible microbial signatures is important for improving our understanding of disease mechanisms and for developing reliable microbiome-based biomarkers in the future.

Based on these findings, we hypothesized that combining multiple independent gut microbiome datasets would help identify microbial changes that are consistently associated with Multiple Sclerosis despite differences between study populations. Rather than focusing on findings from a single cohort, this study investigated whether common alterations in microbial diversity, community composition, and disease-associated bacterial taxa could be identified across independent datasets. To test this hypothesis, three publicly available gut microbiome datasets from clinically characterized MS patients and healthy controls were integrated using a standardized analytical framework. We then evaluated microbial diversity, community composition, phylogenetic relationships, and differential bacterial abundance to identify conserved microbial signatures associated with MS. By identifying reproducible microbial changes linked to immune dysregulation and the loss of beneficial short-chain fatty acid-producing bacteria, this study provides further evidence supporting the microbiota–gut– brain axis in MS and offers a stronger foundation for future microbiome-based biomarkers and therapeutic strategies.

## Materials & Methods

This study employed an integrative cross-cohort observational design to investigate gut microbial alterations associated with Multiple Sclerosis (MS). Three independent human gut microbiome datasets generated using Illumina 16S rRNA amplicon sequencing were systematically analyzed using a standardized analytical workflow to minimize technical variability across datasets. The workflow included sequence quality assessment, denoising, taxonomic classification, phylogenetic reconstruction, microbial diversity analyses, and differential abundance testing. Integrating multiple independent cohorts enabled identification of reproducible microbial signatures associated with disease status rather than cohort-specific observations.

## Data Collection and Dataset Selection

Raw paired-end 16s rRNA amplicon sequence reads using the Illumina NovaSeq 6000 sequencing platform in this study were obtained from three publicly available human gut microbiome BioProjects (PRJEB34168, PRJNA889427, and PRJNA1000059) available through the National Center for Biotechnology Information (NCBI) Sequence Read Archive (SRA) and BioProject databases. These datasets included gut microbiome samples from clinically diagnosed Multiple Sclerosis (MS) patients and healthy control individuals.

Datasets were selected using predefined inclusion criteria to ensure reliable comparison across independent cohorts. Selection was based on the availability of raw FASTQ files, clinically characterized MS and healthy control samples, sufficient sample size, complete sample metadata, and compatibility with a standardized downstream analysis workflow. The inclusion of datasets from different study populations increased the diversity of the analyzed cohorts and reduced the possibility of cohort-specific bias. This integrative approach improved the robustness of the analysis and enabled the identification of microbial signatures that were consistently associated with MS across independent studies.

After data collection, all raw FASTQ files were imported into QIIME2 (version 2023.2) and converted into QIIME2 artifact (.qza) format to ensure standardized processing throughout the analytical workflow. All analyses were performed using QIIME2 version 2023.2 through the Galaxy Europe platform (https://usegalaxy.eu/), which provides a reproducible cloud-based computational environment for microbiome data analysis.

## Demultiplexing and Denoising of Sequences

Before sequence processing, raw FASTQ files were evaluated using FastQC (version 0.11.9) to assess sequence quality, GC-content distribution, sequence length distribution, adapter contamination, and per-base quality scores. Quality reports were reviewed to determine appropriate trimming positions before denoising. Raw sequencing reads were first demultiplexed to assign each sequence to its corresponding biological sample based on the barcode information generated during library preparation. Sequence quality profiles were then examined to determine suitable trimming positions and to retain high-quality reads for downstream analysis [19]. Low-quality regions at the ends of reads were removed to reduce sequencing artifacts and improve the overall quality of the dataset.

Denoising was performed using the DADA2 plugin implemented in QIIME2 version 2023.2. Quality filtering parameters were selected after inspection of sequence quality profiles generated by FastQC. Forward reads were truncated at 249 bp, reverse reads at 220 bp, while no bases were removed from the 5′ end of either read (trim-left = 0 bp). The maximum expected error threshold (maxEE) was set to 2.0 for both forward and reverse reads. A minimum overlap of 12 bp was required for paired-end read merging. Samples were processed using the independent pooling strategy, and PCR chimeras were identified and removed using the consensus method. These parameters were selected to retain high-quality sequence regions while minimizing sequencing errors and PCR artefacts. Truncation positions were selected based on per-base quality score profiles. Forward reads maintained Phred quality scores above Q30 for approximately the first 240 bp, whereas reverse read quality decreased after approximately 200 bp. The selected truncation lengths preserved high-quality sequence information while maintaining sufficient overlap for accurate paired-end merging.

## Taxonomy Classification and Phylogenetic Analysis

Taxonomic classification was performed to identify the microbial composition of each sample and to compare the distribution of bacterial taxa between Multiple Sclerosis (MS) patients and healthy controls. Representative 16S rRNA amplicon sequence variants (ASVs) were assigned to taxonomic groups ranging from phylum to genus level using the QIIME2 Feature Classifier plugin with a pre-trained Naive Bayes classifier based on the SILVA reference database. This standardized reference database enabled consistent and reliable taxonomic assignment across all datasets.

To investigate the evolutionary relationships among the identified microorganisms, a phylogenetic tree was constructed using the q2-phylogeny pipeline in QIIME2, which incorporates sequence alignment with MAFFT and tree construction using FastTree [20,21]. The resulting phylogenetic tree was used in downstream diversity analyses by incorporating evolutionary relationships among microbial taxa. This approach provided additional biological context for interpreting differences in gut microbial community structure between MS patients and healthy controls and supported a more comprehensive assessment of microbial diversity.

## Diversity Analysis

Microbial diversity was assessed to compare the composition and structure of gut microbial communities between Multiple Sclerosis (MS) patients and healthy controls using the QIIME2 analytical framework on the Galaxy Europe platform. Both alpha and beta diversity analyses were performed to evaluate microbial diversity within and between study groups.

Alpha diversity was used to measure the richness and evenness of microbial communities within individual samples. Rarefaction analysis was performed before diversity estimation to identify an appropriate sequencing depth that maximized sample retention while minimizing sampling bias. The selected rarefaction depth was subsequently applied uniformly to all diversity analyses [22]. Core alpha diversity metrics, including Observed Features, Shannon diversity index, Simpson diversity index, and Faith’s Phylogenetic Diversity (PD), were then calculated to provide both non-phylogenetic and phylogenetic measures of microbial diversity [23,24]. These metrics allowed a comprehensive assessment of microbial richness, evenness, and evolutionary diversity across all samples.

Beta diversity was used to examine differences in microbial community composition between MS patients and healthy controls. Distance matrices based on Bray-Curtis, Jaccard, Weighted UniFrac, and Unweighted UniFrac metrics were generated to evaluate similarities and differences in microbial communities while considering both taxonomic composition and phylogenetic relationships. Together, alpha and beta diversity analyses provided a comprehensive evaluation of gut microbial diversity and community structure across the study cohorts.

## Statistical Significance Testing of Group Differences

Statistical analyses were performed using QIIME2 plugins implemented through the Galaxy Europe platform. Differences in alpha diversity between MS patients and healthy controls were evaluated using the Kruskal-Wallis test. Differences in microbial community composition (beta diversity) were assessed using PERMANOVA (ADONIS) with 999 permutations based on Bray-Curtis, Jaccard, Weighted UniFrac, and Unweighted UniFrac distance matrices [25]. For all statistical analyses, a p-value of less than 0.05 was considered statistically significant. The R² value obtained from PERMANOVA was used to estimate the proportion of variation explained by disease status, allowing evaluation of whether gut microbial community composition differed significantly between MS patients and healthy controls [26]. Before differential abundance testing, the feature table was transformed using the centered log-ratio (CLR) transformation to account for the compositional nature of microbiome sequencing data. CLR transformation minimizes biases introduced by relative abundance data and improves statistical interpretation of microbial differences between study groups.

## Differential Abundance Analysis

Differential abundance analysis was performed using the Analysis of Composition of Microbiomes (ANCOM) method within the QIIME2 environment to identify bacterial taxa that differed significantly between Multiple Sclerosis (MS) patients and healthy controls [27,28]. ANCOM accounts for the compositional nature of microbiome sequencing data and enables reliable identification of differentially abundant taxa while reducing false-positive results. Bacterial taxa showing significant differences between study groups were considered potential microbial signatures associated with MS and were used to better understand disease-associated alterations in the gut microbiome. Differential abundance results were visualized using volcano plots, where CLR-transformed abundance values were plotted against ANCOM W-statistics to facilitate identification of significantly enriched or depleted microbial taxa. Scatter plots were generated to visualize relationships between CLR-transformed abundances and ANCOM W-statistics for differentially abundant microbial taxa.

## Taxonomic Feature Visualization

Taxonomic profiles generated during classification were visualized using bar plots and heatmaps to facilitate comparison of microbial communities between MS patients and healthy controls. Relative abundance bar plots were used to illustrate the distribution of bacterial taxa across individual samples and study groups. Taxa were collapsed at the genus level before visualization to provide a clearer representation of microbial composition. Heatmaps were generated to display the relative abundance of bacterial taxa across samples, allowing identification of shared microbial patterns as well as taxa that were enriched or depleted in the disease group. These visualizations supported the interpretation of microbial community structure and assisted in identifying potential disease-associated microbial signatures.

## Quality Control and Data Validation

Multiple quality control steps were incorporated throughout the analytical workflow to ensure reliable and reproducible results. Sequence quality was assessed before denoising, and low-quality reads and chimeric sequences were removed during data processing. Read retention statistics were examined after denoising to confirm that sufficient high-quality sequences were retained for downstream analyses. Taxonomic assignments were reviewed for consistency across all datasets, and rarefaction curves were evaluated to verify adequate sequencing depth. Diversity and phylogenetic analyses were performed only after quality-filtered, high-confidence amplicon sequence variants (ASVs) had been obtained. These quality assessment procedures ensured that all downstream analyses were based on reliable sequencing data.

## Results & Discussion

### Study Cohort and Dataset Characteristics

A total of 234 human stool samples from three independent publicly available gut microbiome studies were included in this analysis. The combined dataset consisted of 137 clinically diagnosed Multiple Sclerosis (MS) patients and 97 healthy control individuals. The selected datasets (PRJEB34168, PRJNA889427, and PRJNA1000059) were generated using Illumina-based 16S rRNA amplicon sequencing and were chosen because they provided raw sequencing data together with complete clinical metadata, allowing consistent analysis across independent cohorts. The three studies represented different age groups and included both male and female participants, providing a diverse study population for evaluating gut microbiome alterations associated with MS. PRJEB34168 included adult male and female participants between 21 and 66 years of age, PRJNA889427 consisted of middle-aged adults, whereas PRJNA1000059 represented a younger female cohort. Integrating these independent cohorts reduced the influence of study-specific variation and increased the robustness of the biological observations. The demographic characteristics of all included datasets are summarized in Table 1, while the distribution of patient groups, gender, and age is presented in Figure 2.

**Figure 1.**
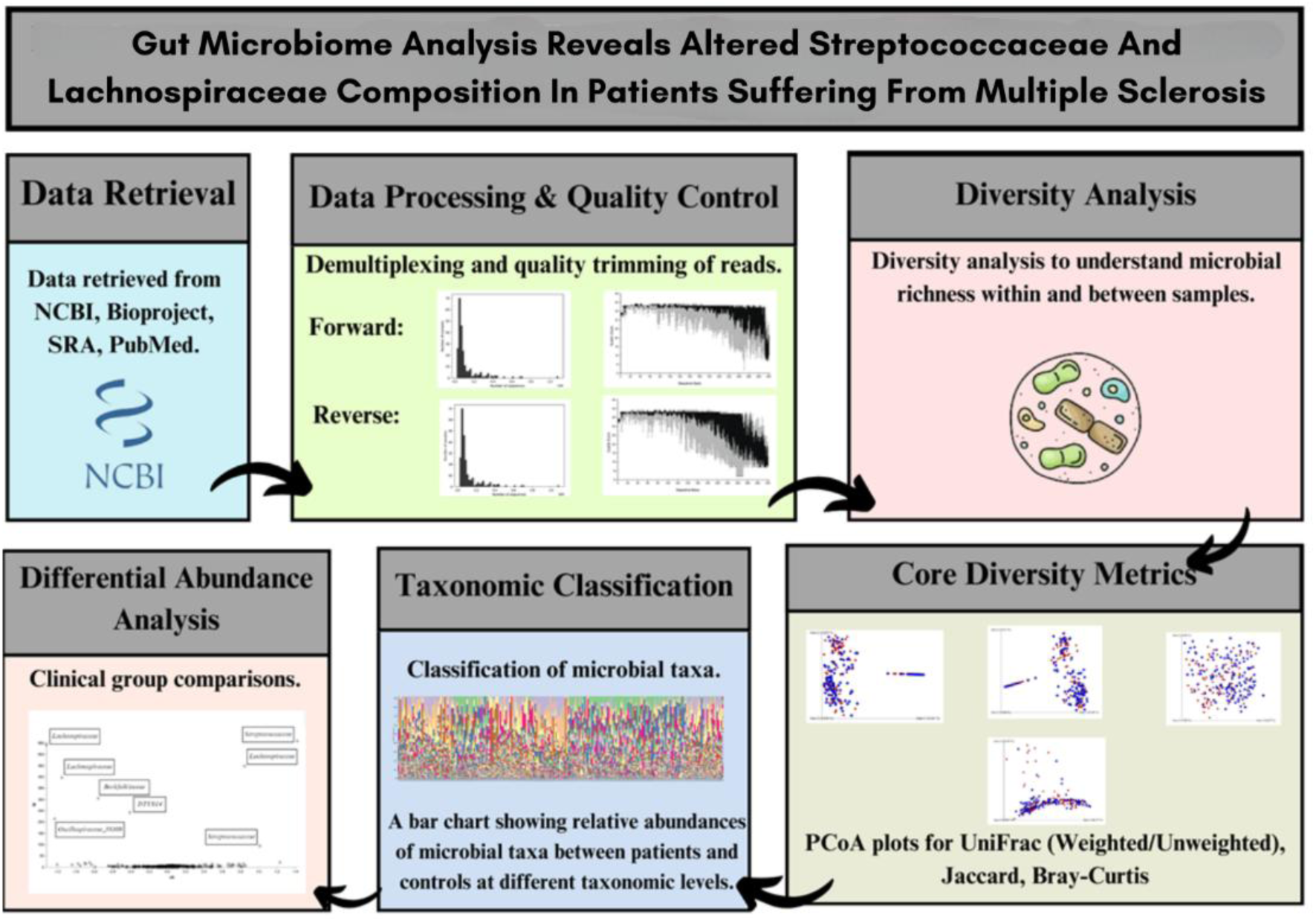
Overall experimental design illustrating dataset integration, standardized microbiome processing, taxonomic characterization, diversity profiling, statistical analyses, and identification of disease-associated microbial signatures in Multiple Sclerosis.

**Figure 2.**
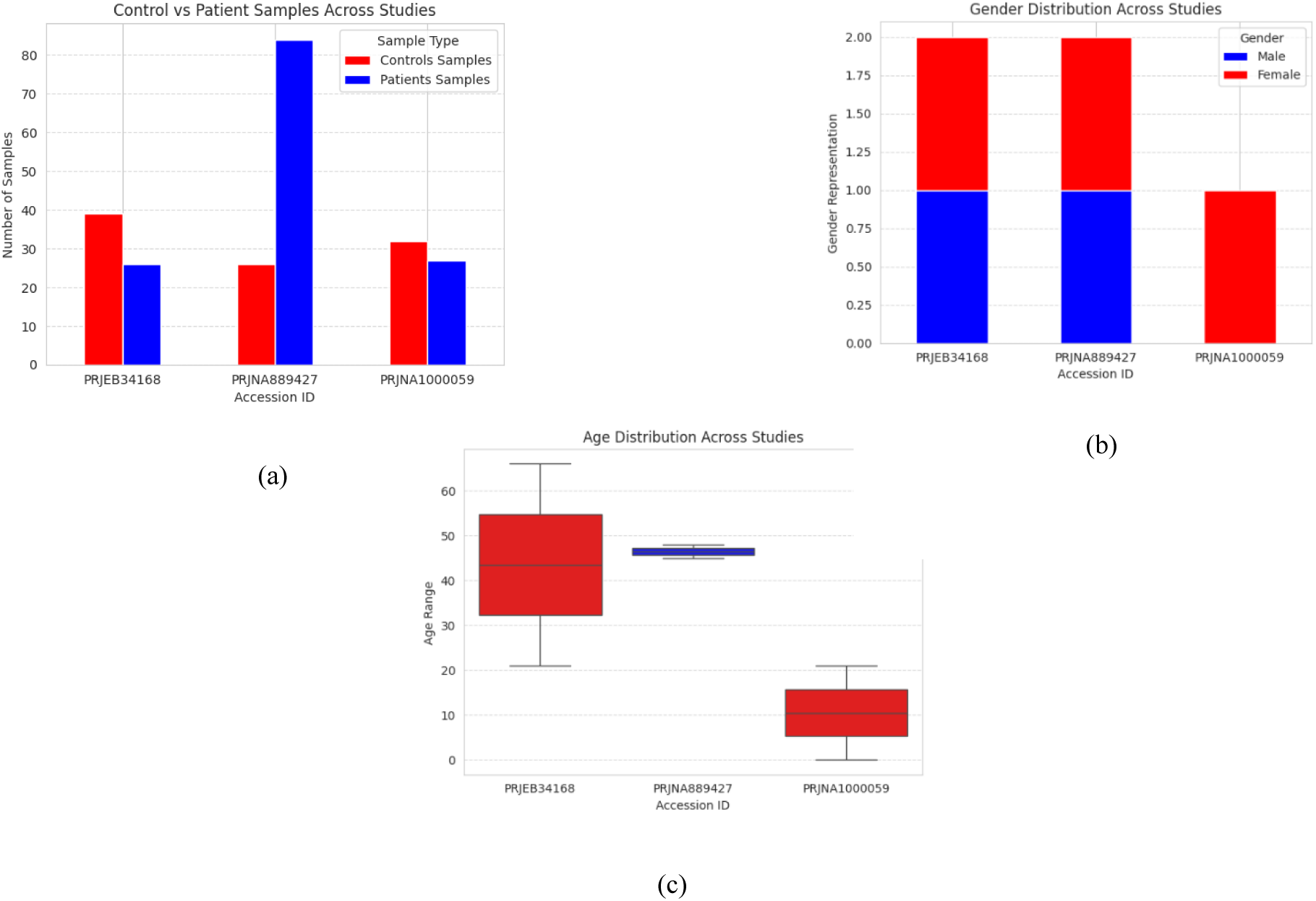
Demographic characteristics of the integrated study population. (a) Distribution of Multiple Sclerosis patients and healthy controls, (b) gender distribution across the included cohorts, and (c) age distribution of study participants. The inclusion of independent cohorts with diverse demographic characteristics improved the robustness of the microbiome analysis and reduced cohort-specific bias.

**Table 1.** Study Participants and Sample Characteristics

| BioProject Accession | Sample Type | Gender | Controls Samples | Patients Samples | Age Range (in years) | Geographic Region | Ref |
| --- | --- | --- | --- | --- | --- | --- | --- |
| PRJEB34168 | Stool | Female; Male | 39 | 26 | 21-66 | USA | [29] |
| PRJNA889427 | Stool | Female; Male | 26 | 84 | 45- 48 | USA | [30] |
| PRJNA1000059 | Stool | Female | 32 | 27 | < 21 | USA | [31] |

### High-Quality Sequencing Data Support Reliable Microbiome Profiling

Raw sequencing reads from all 234 samples were successfully processed through a standardized quality control workflow before downstream microbiome analysis. Initial quality assessment showed that the sequencing data were of sufficient quality for reliable processing. Sequence quality profiles were used to identify appropriate trimming positions, allowing low-quality regions to be removed while retaining high-quality reads for subsequent analyses.

Read count distributions demonstrated that most samples contained adequate sequencing depth for microbiome profiling, although some variation in sequencing depth was observed among samples (Figure 3). Rarefaction depth was selected based on these distributions to ensure fair comparison of microbial diversity across all samples.

**Figure 3.**
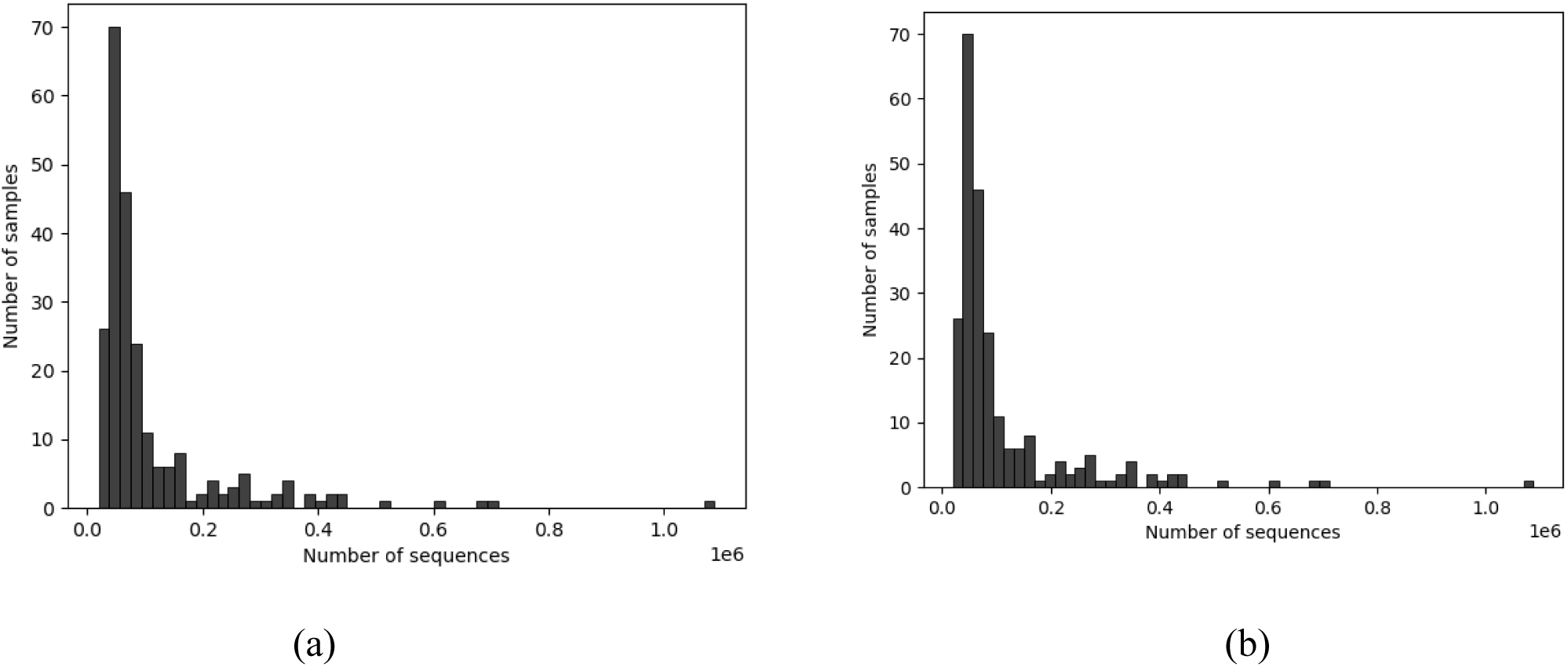
Read Distribution Histograms forward reads (a) Forward read count distribution across all samples (b) Reverse read count distribution across all samples. Most samples had read counts below 200,000, with some exceeding 1 million. This informed the rarefaction depth used in diversity analysis.

Following quality filtering and denoising, sequencing errors and chimeric sequences were removed, resulting in a high-confidence set of amplicon sequence variants (ASVs) for downstream taxonomic and diversity analyses. Overall, the quality control process confirmed that the sequencing data were suitable for reliable characterization of gut microbial communities in both MS patients and healthy controls.

### Gut Microbial Diversity Is Reduced in Multiple Sclerosis

To determine whether gut microbial diversity differs between Multiple Sclerosis (MS) patients and healthy controls, alpha diversity was evaluated using the Shannon diversity index following standardized rarefaction analysis. Alpha diversity reflects both the richness and evenness of microbial communities and provides an overall measure of gut microbial health. Reduced alpha diversity is commonly considered an indicator of microbial imbalance and has been associated with several immune-mediated disorders. The alpha rarefaction curves reached a stable plateau at approximately 1,100 sequencing reads, indicating that the sequencing depth was sufficient to capture most microbial diversity present in each sample (Figure 5). Based on these results, a rarefaction depth of 10,275 reads was selected to ensure consistent comparison while retaining the maximum number of samples for downstream analyses. Comparison of Shannon diversity between study groups showed that MS patients had significantly lower microbial diversity than healthy controls (Kruskal-Wallis test, p = 0.024). This reduction suggests that the gut microbiome of MS patients is less diverse and may have lost beneficial microbial members that normally contribute to intestinal homeostasis and immune regulation. Similar reductions in microbial diversity have been reported in previous studies of autoimmune and neuroinflammatory diseases, supporting the presence of gut microbial dysbiosis in MS. Although reduced diversity alone cannot explain disease development, it provides evidence of an altered gut microbial ecosystem that may contribute to disease progression through disruption of normal host-microbiota interactions.

**Figure 4.**
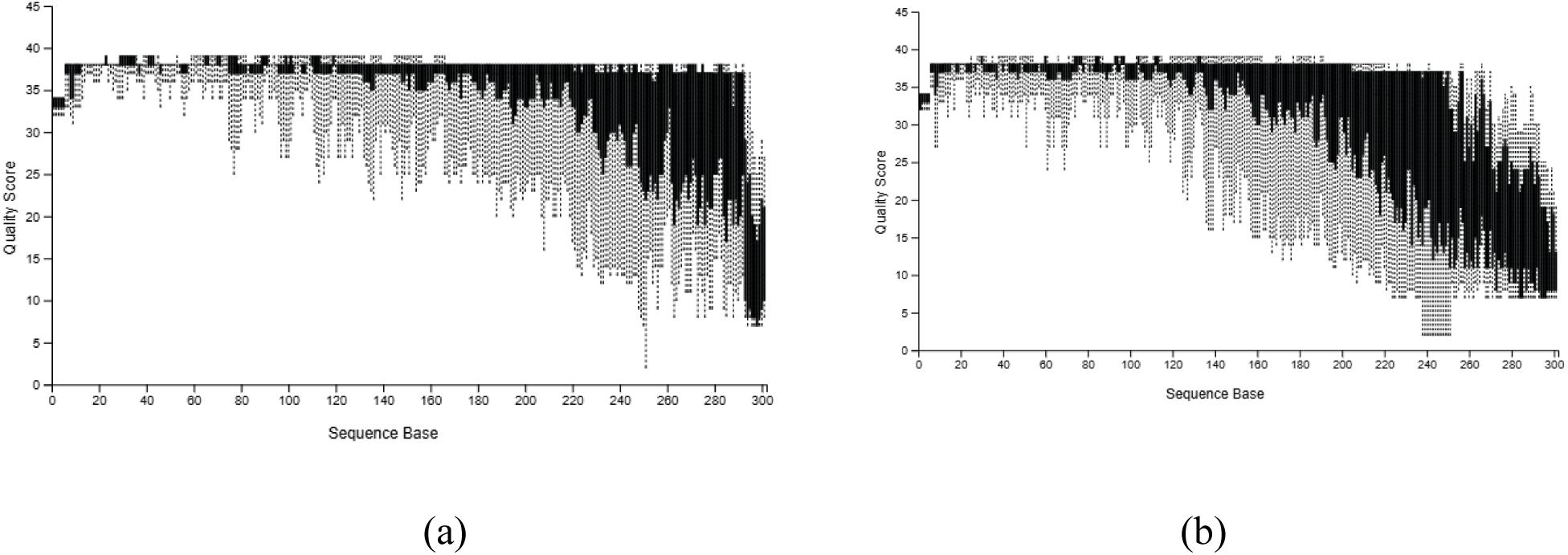
Quality Score Distribution Plots (a) Quality score profiles for forward reads across base position. (b) Quality score profiles for reverse reads. Forward reads retained high Phred scores (>Q30) up to ∼240 bp; reverse reads dropped in quality after ∼200 bp. Truncation was set at 249 bp (forward) and 220 bp (reverse) to preserve high-quality regions.

**Figure 5:**
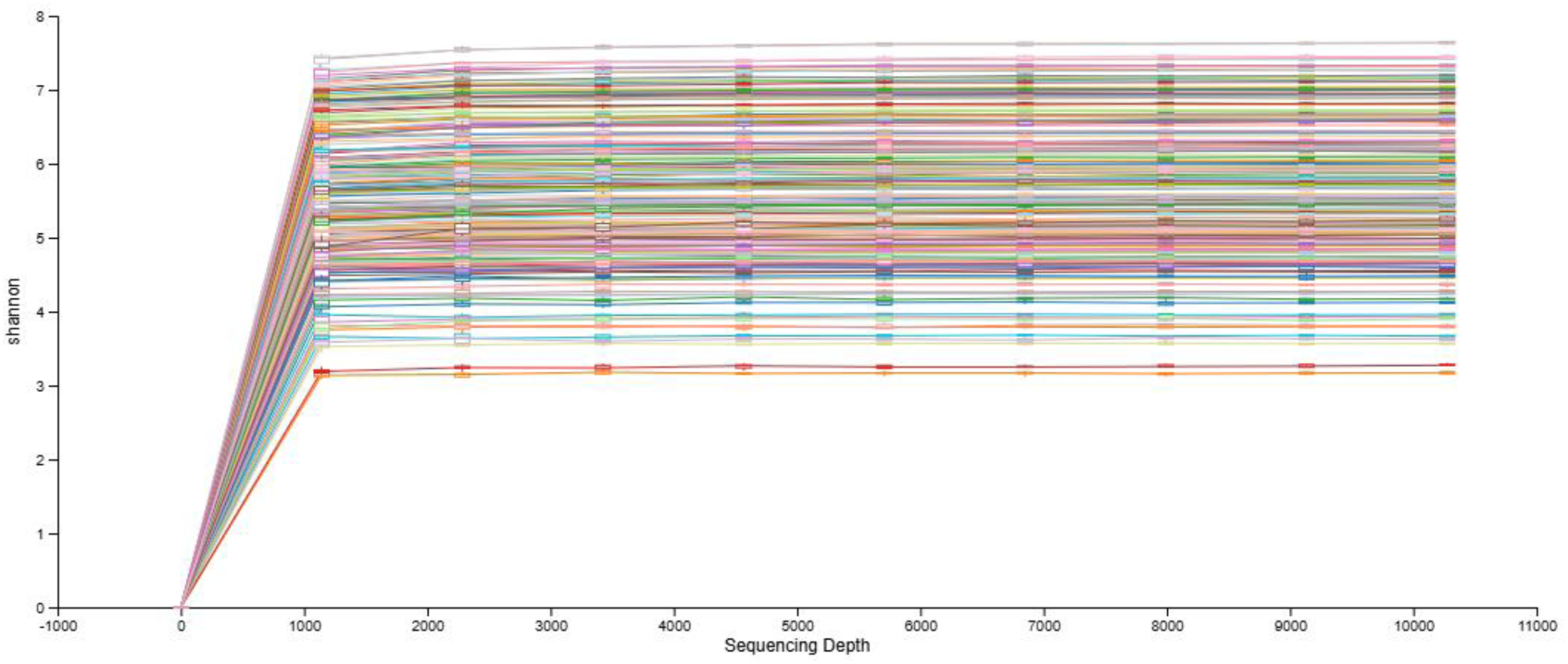
Alpha rarefaction curves showing Shannon diversity across increasing sequencing depths for each sample. Most curves plateau around 1,100 reads, indicating sufficient sampling depth. A rarefaction threshold of 10,275 reads was selected for downstream analysis to ensure both sample retention and analytical consistency.

### Multiple Sclerosis Reshapes Gut Microbial Community Structure

To investigate whether the overall composition of the gut microbiome differed between Multiple Sclerosis (MS) patients and healthy controls, beta diversity was evaluated using Bray-Curtis, Weighted UniFrac, Unweighted UniFrac, and Jaccard distance metrics. Although these metrics assess microbial community differences from different perspectives, all four analyses consistently demonstrated separation between MS and healthy control samples, indicating that disease status is associated with changes in gut microbial community structure.

Principal Coordinate Analysis (PCoA) based on Bray-Curtis distances showed clear clustering of MS patients and healthy controls, demonstrating differences in microbial composition based on taxon abundance (Figure 6). Similar clustering patterns were observed using both Weighted and Unweighted UniFrac distances (Figures 7.1 and 7.2), indicating that differences were associated not only with microbial abundance but also with the evolutionary relationships among bacterial taxa. The Jaccard analysis also demonstrated separation between study groups (Figure 7.3), suggesting that the presence or absence of specific bacterial taxa contributed to the observed microbial differences. Although partial overlap between some samples was observed, the overall clustering pattern remained consistent across all distance metrics.

**Figure 6.**
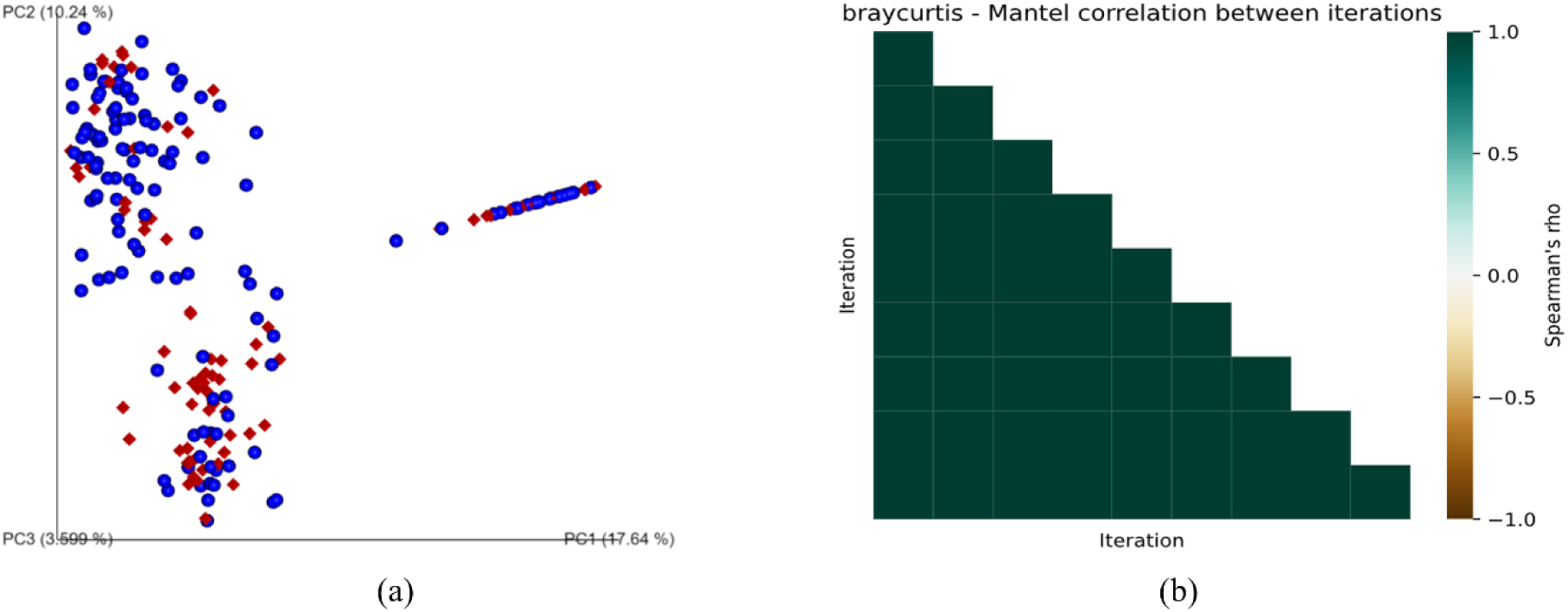
(a) Emperor jackknifed PCoA plot based on Bray-Curtis dissimilarities, showing clustering of MS patient samples (blue spheres) and control samples (red diamond). (b) Heatmap showing Bray-Curtis dissimilarity matrix, illustrating correlations between iterations of diversity metrics across rarefaction trials.

**Figure 7.**
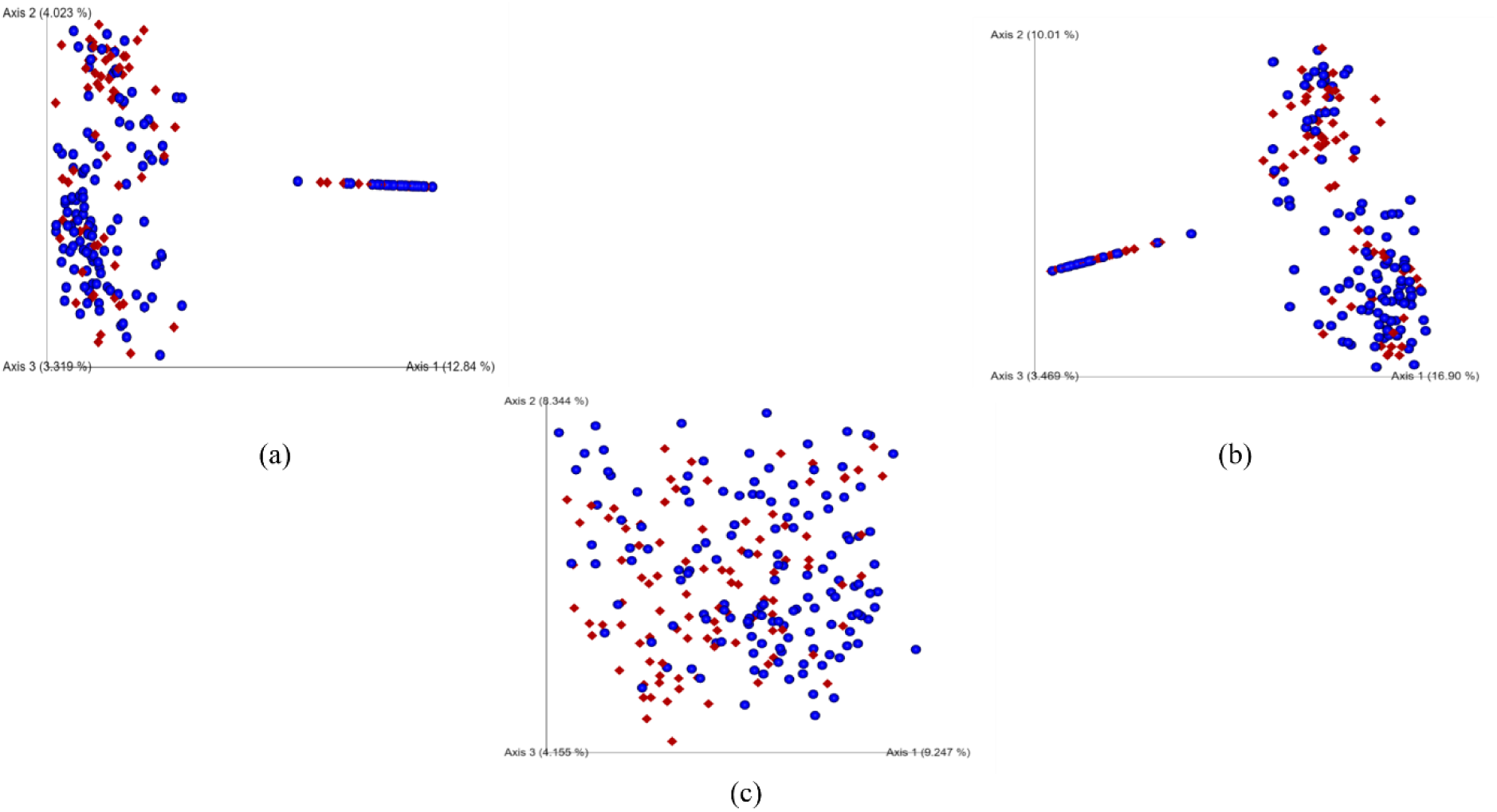
Principal Coordinate Analysis (PCoA) of gut microbial communities in Multiple Sclerosis MS patients (blue spheres) and controls (red diamonds) using different beta diversity distance metrics. (a) Unweighted UniFrac PCoA showing separation between MS and healthy control samples based on the presence or absence of bacterial taxa, indicating differences in microbial community membership. (b) Weighted UniFrac PCoA demonstrating differences between MS patients and healthy controls after incorporating both bacterial abundance and phylogenetic relationships, supporting disease-associated changes in microbial community structure. (c) Jaccard PCoA illustrating variation in microbial community composition based on the presence or absence of bacterial taxa. The clustering patterns observed across all three-distance metrics consistently indicate alterations in the gut microbiome associated with Multiple Sclerosis.

Statistical analysis further supported these observations. PERMANOVA analysis showed significant differences in microbial community composition between MS patients and healthy controls across all beta diversity metrics (Bray-Curtis, p = 0.001; Weighted UniFrac, p = 0.001; Unweighted UniFrac, p = 0.002; Jaccard, p = 0.001). ADONIS analysis also confirmed significant group separation (p = 0.004), while the corresponding PERMANOVA result (R² = 0.12, p = 0.001) indicated that disease status explained a measurable proportion of the observed variation in gut microbial composition.

Together, these findings demonstrate that the gut microbiome of MS patients differs consistently from that of healthy individuals. The agreement among four independent beta diversity metrics strengthens the reliability of these observations and suggests that microbial alterations in MS involve changes in both bacterial composition and phylogenetic structure. Although some variation among individual samples is expected because of biological diversity, the consistent clustering observed across three independent cohorts increases confidence that these microbial patterns are associated with Multiple Sclerosis rather than being specific to a single study population.

ADONIS test results confirmed statistically significant differences in beta diversity between Multiple Sclerosis (MS) patients and healthy control groups. The analysis yielded a p-value of 0.004, indicating that the observed dissimilarities in microbial community composition were unlikely due to chance. The corresponding PERMANOVA analysis supported these findings, with a p-value = 0.001 and R² = 0.12, further validating the distinct clustering observed in the Principal Coordinates Analysis (PCoA) plots (Figure 6). These results suggest that the gut microbiome composition in MS patients is significantly different from that of healthy individuals. These findings reinforce the hypothesis that MS is associated with distinct alterations in the gut microbial ecosystem, which may play a role in disease onset or progression.

### Conserved Microbial Dysbiosis Characterizes Multiple Sclerosis

To identify microbial changes consistently associated with Multiple Sclerosis (MS), the gut microbiome composition of MS patients and healthy controls was compared at both the family and genus levels. Taxonomic profiling revealed a consistent pattern of microbial dysbiosis across the integrated datasets, characterized by an increase in potentially pro-inflammatory bacterial groups together with a reduction in beneficial bacteria involved in maintaining intestinal homeostasis.

At the family level, *Streptococcaceae* and *Bacteroidaceae* showed higher relative abundance in MS patients, whereas *Lachnospiraceae* and *Ruminococcaceae* were consistently reduced compared with healthy controls (Table 2). The depletion of *Lachnospiraceae* and *Ruminococcaceae* is particularly important because these bacterial families include many short-chain fatty acid (SCFA)-producing microorganisms that help maintain intestinal barrier integrity and regulate immune responses. Their reduced abundance suggests a loss of beneficial microbial functions that may contribute to the inflammatory environment observed in MS.

**Table 2.** Differentially Abundant Bacterial Families Between MS and Controls

| Bacterial Family | Change in MS | Potential Biological Relevance |
| --- | --- | --- |
| *Streptococcaceae* | Increased | Associated with pro-inflammatory microbial communities |
| *Bacteroidaceae* | Increased | Altered gut microbial composition in MS |
| *Lachnospiraceae* | Decreased | Major producer of short-chain fatty acids (SCFAs) |
| *Ruminococcaceae* | Decreased | Supports gut barrier integrity and immune regulation |

A similar pattern was observed at the genus level (Table 3 and Figure 8). Streptococcus showed higher relative abundance in MS patients, while beneficial genera such as Blautia and *Ruminococcus* were reduced. Both Blautia and Ruminococcus are recognized as important producers of SCFAs, including butyrate, which supports gut barrier function and helps regulate immune homeostasis. Their depletion may therefore reduce anti-inflammatory microbial activity and contribute to immune dysregulation associated with Multiple Sclerosis.

**Figure 8.**
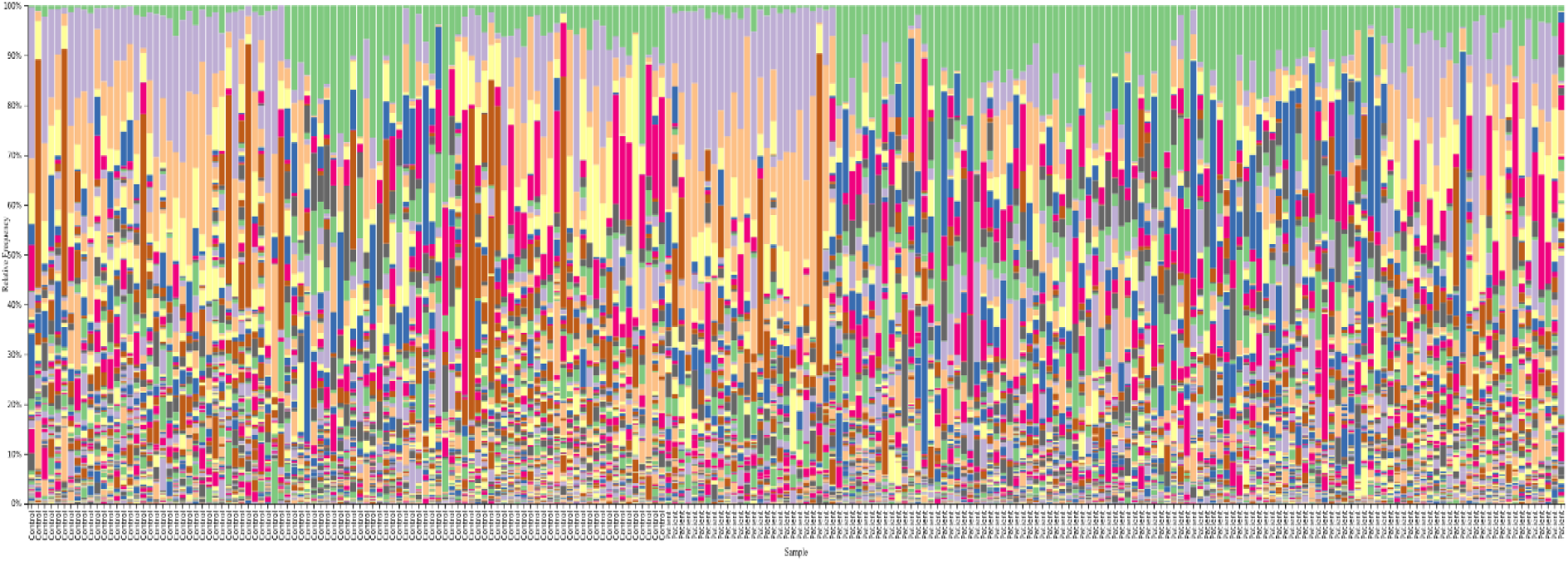
Bar plot of taxonomic composition at the genus level, highlighting the distribution of dominant bacterial genera in MS patient groups (blue) and healthy controls (red). This comparison visualizes key compositional shifts contributing to the dysbiotic gut profile in MS.

**Table 3.** Genus-Level Differences Observed Between MS Patients and Controls

| **Bacterial Genus** | **Change in MS** | **Potential Biological Relevance** |
| --- | --- | --- |
| *Streptococcus* | Increased | Associated with inflammatory responses |
| *Blautia* | Decreased | SCFA-producing beneficial bacterium |
| *Ruminococcus* | Decreased | Contributes to butyrate production and gut health |

Overall, these findings demonstrate a conserved pattern of gut microbial dysbiosis characterized by enrichment of *Streptococcaceae*-related taxa together with depletion of SCFA-producing members of the *Lachnospiraceae* and *Ruminococcaceae* families. The consistent observation of these microbial changes across three independent cohorts strengthens the evidence that disruption of beneficial gut bacteria is closely associated with MS and supports the role of the microbiota-gut-brain axis in disease pathogenesis.

### Differential Abundance Identifies Immune-Associated Microbial Signatures

To identify bacterial taxa that consistently differed between Multiple Sclerosis (MS) patients and healthy controls, differential abundance analysis was performed at the genus level using ANCOM. The analysis identified several microbial taxa that showed significant and reproducible differences between the two groups (Figure 9 and Table 4). Among all detected genera, Streptococcus showed the strongest association with MS (W = 559), indicating that it was consistently enriched across the patient samples. In contrast, beneficial bacteria including *Faecalibacterium* (W = 398) and unclassified members of the *Lachnospiraceae* family (W = 542) were reduced in the MS group. Other genera, such as *Clostridium*_Q, also showed lower abundance in MS, whereas *Blautia*, *Borkfalkia*, DTU014, *Limivicinus*, and *Lactococcus*_A did not reach statistical significance in the present analysis (Table 4)

**Figure 9.**
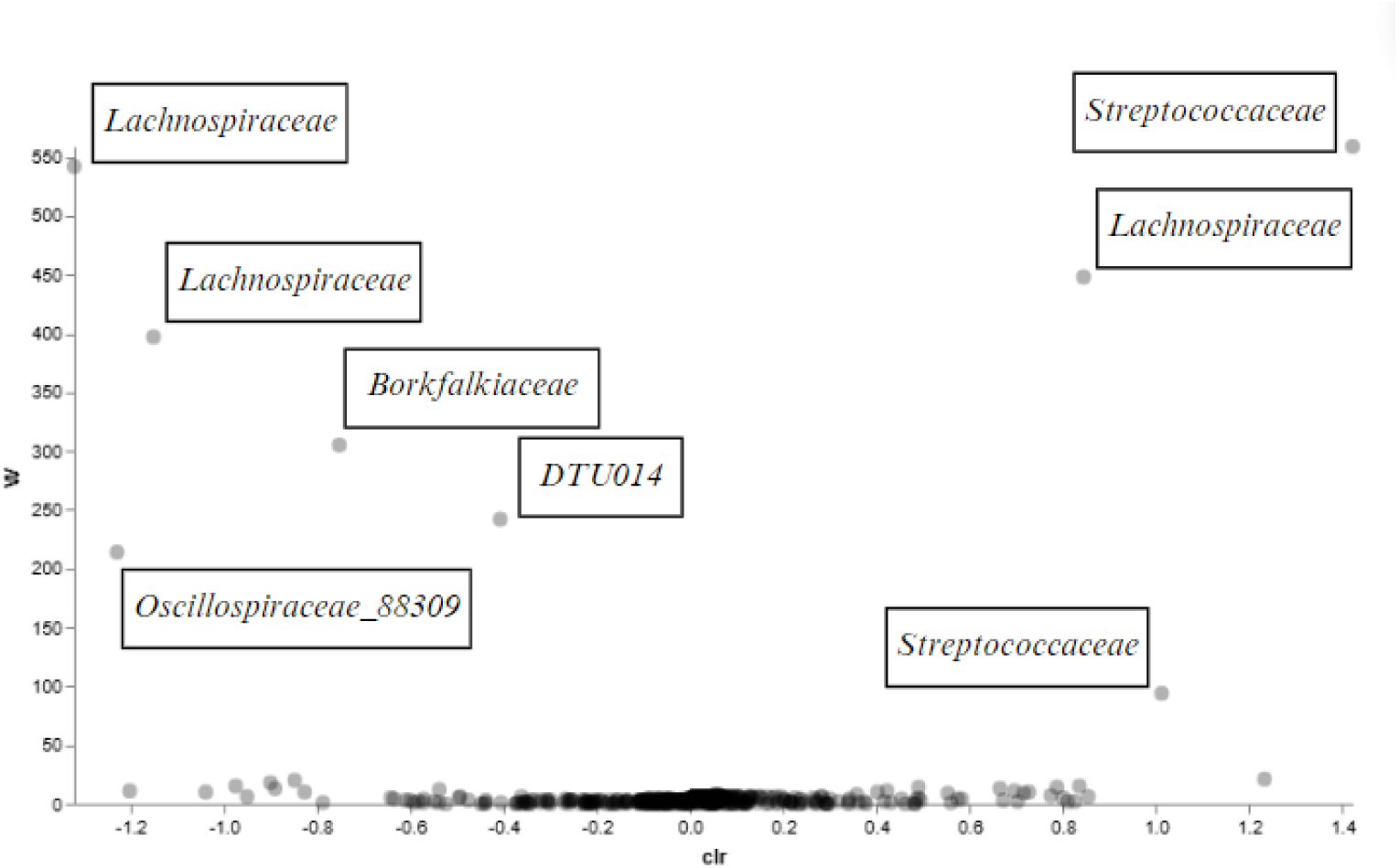
illustrates the differentially abundant taxa identified in this study. Taxa with higher W-statistics showed more consistent differences between the MS and healthy control groups, indicating stronger evidence of disease association. The results demonstrate that enrichment of Streptococcus and depletion of SCFA-producing bacteria represent the most reproducible microbial signatures observed across the three independent cohorts.

**Table 4.**
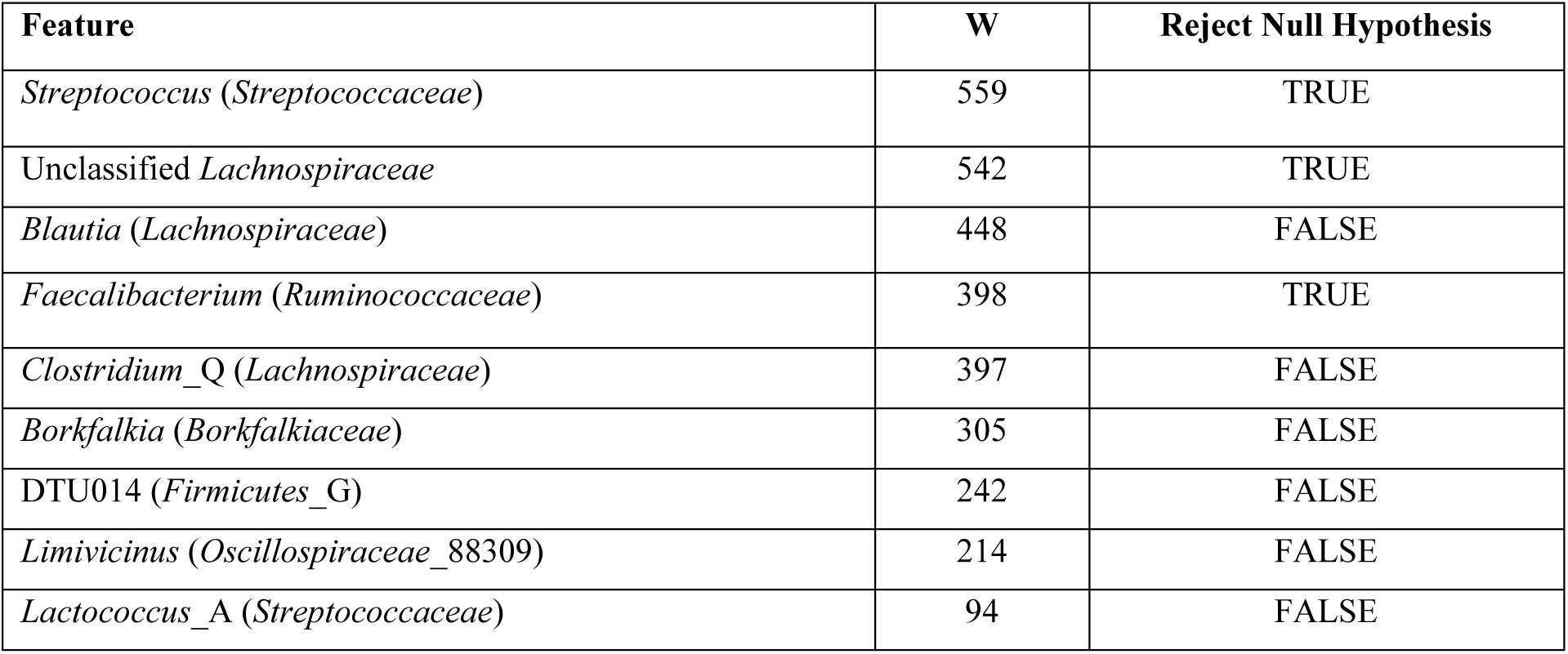
Differentially abundant bacterial genera identified between Multiple Sclerosis patients and healthy controls..

These findings indicate that MS is associated with a reproducible shift in gut microbial composition, characterized by an increase in potentially pro-inflammatory bacteria together with a reduction in beneficial short-chain fatty acid (SCFA)-producing microorganisms.

Similar microbial patterns have been reported in previous studies, supporting the idea that gut microbial imbalance is closely associated with MS development and progression [33].

To further examine these microbial differences, the significant taxa identified by ANCOM were visualized using a scatter plot (Figure 10). Taxa with positive abundance values, including Streptococcus, were more common in MS patients, whereas *Faecalibacterium*, *Blautia*, and members of the *Lachnospiraceae* family showed higher abundance in healthy controls. These patterns agreed with the taxonomic analyses presented earlier and further confirmed that the gut microbiome differs consistently between the two study groups.

**Figure 10.**
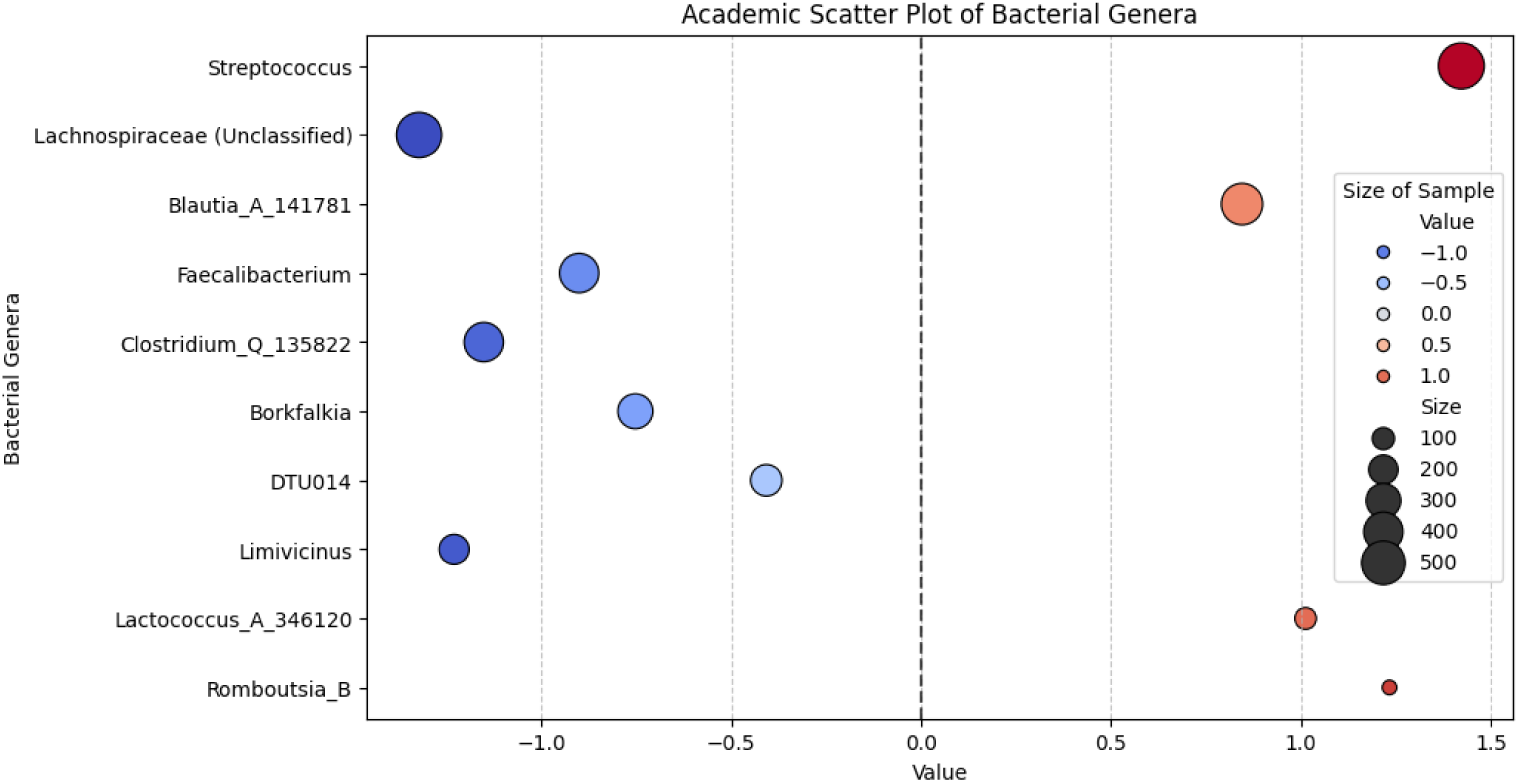
Scatter plot showing differential abundance of key bacterial genera associated with Multiple Sclerosis. Disease-associated taxa, particularly Streptococcus, showed increased abundance, whereas beneficial SCFA-producing genera, including Faecalibacterium, Blautia, and Lachnospiraceae, were reduced in MS patients.

The overall findings of this study suggest that gut microbial dysbiosis is closely associated with Multiple Sclerosis. Across three independent cohorts, the most consistent pattern was the enrichment of Streptococcus together with the depletion of *Lachnospiraceae*, *Ruminococcaceae*, and *Faecalibacterium*. These bacterial groups are known to influence immune regulation and intestinal homeostasis. Previous studies have also reported increased abundance of Streptococcus in MS and other neuroinflammatory disorders, suggesting that this genus may promote inflammatory immune responses through interactions with host immunity [33–35]. In contrast, members of the *Lachnospiraceae* and *Ruminococcaceae* families are important producers of short-chain fatty acids, especially butyrate, which supports gut barrier integrity and helps regulate immune function [36–38]. Therefore, their depletion may reduce anti-inflammatory activity and contribute to immune imbalance in MS.

Our observations are consistent with earlier microbiome studies reporting reduced SCFA-producing bacteria and increased pro-inflammatory taxa in MS patients [33–39]. By combining three independent publicly available cohorts and processing all samples using the same analytical workflow, this study identified microbial changes that were consistently observed across different datasets rather than being limited to a single study. This strengthens confidence that the identified microbial signatures are associated with disease status instead of cohort-specific variation.

These findings also have potential clinical importance. Although the present results do not establish a causal relationship, the consistent enrichment of *Streptococcus* and depletion of beneficial SCFA-producing bacteria suggest that these microbial groups may serve as candidate biomarkers for MS. They may also represent potential targets for future microbiome-based interventions, including dietary modification, prebiotics, probiotics, or other approaches aimed at restoring gut microbial balance. However, these applications require further validation in well-designed clinical studies.

## Conclusion

This study provides critical insights into the gut microbiome alterations associated with Multiple Sclerosis (MS), revealing a distinct microbial signature that may contribute to disease progression. A significant increase in *Streptococcus* abundance was observed in MS patients, accompanied by a marked depletion of *Lachnospiraceae* and *Faecalibacterium* (*Ruminococcaceae*) are key microbial groups known for their roles in maintaining gut and immune homeostasis. The loss of beneficial short-chain fatty acid (SCFA)-producing bacteria, particularly *Lachnospiraceae* and *Faecalibacterium*, may weaken anti-inflammatory defenses, potentially intensifying immune dysregulation. At the same time, the expansion of *Streptococcus*, a genus linked to inflammatory pathways, suggests a possible interaction between gut microbiota and neuroinflammation, reinforcing the intricate relationship between the gut and the central nervous system. While the findings strongly indicate a link between gut microbial imbalances and MS, further investigations will be valuable in uncovering the precise biological mechanisms underlying these changes. Expanding research efforts through functional microbiome studies, cross-cohort validations, and longitudinal analyses will enhance our ability to translate these observations into actionable clinical insights. Moreover, integrating multi-omics approaches, such as metagenomics and metabolomics, will provide a more comprehensive understanding of the microbiome’s role in MS. These advancements will pave the way for the development of microbiome-targeted interventions, including precision probiotics, dietary modifications, and personalized therapeutic strategies aimed at restoring gut microbial balance and potentially alleviating disease symptoms.

By deepening our understanding of the microbiome’s impact on MS pathophysiology, this research not only contributes to the growing field of neuroimmunology but also opens new doors for innovative, gut-centered therapeutic approaches that may improve patient outcomes.

## References

1. B Wootla, M Eriguchi, M Rodriguez. 2012. Is Multiple Sclerosis an Autoimmune Disease?, Autoimmune Dis. 1: 969657.

2. C Matute-Blanch, X Montalban, M Comabella. 2018. Multiple sclerosis, and other demyelinating and autoimmune inflammatory diseases of the central nervous system. Handb Clin. Neurol. 146: 67–84.

3. R Höftberger and H. Lassmann. 2018. Inflammatory demyelinating diseases of the central nervous system. Handb. Clin. Neurol. 145: 263–283.

4. B Sneha, G Lauren, C Langston. 2023. Managing the Symptoms of Multiple Sclerosis. Med. Res. Arch. 11. doi10.18103/mra.v11i2.3539.

5. Laura L Laslett, Cynthia Honan, Jason A Turner, Baye Dagnew, Julie A Campbell, Tiffany K Gill, et al. 2022. Poor sleep and multiple sclerosis: associations with symptoms of multiple sclerosis and quality of life. J. Neurol. Neurosurg. Psychiatry 93: 1162–1165.

6. Yan Zhang, Bruce V Taylor, Steve Simpson Jr, Leigh Blizzard, Julie A Campbell, Andrew J Palmer, et al. 2020. Feelings of depression, pain and walking difficulties have the largest impact on the quality of life of people with multiple sclerosis, irrespective of clinical phenotype. Mult. Scler. 27: 1262–1275.

7. RHB Benedict, MP Amato, J DeLuca, JJG Geurts. 2020. Cognitive impairment in multiple sclerosis: clinical management, MRI, and therapeutic avenues. Lancet Neurol. 19: 860–871.

8. M Margoni, P Preziosa, MA Rocca, M Filippi. 2023. Depressive symptoms, anxiety and cognitive impairment: emerging evidence in multiple sclerosis. Transl. Psychiatry 13: 1–13.

9. HF Harbo, R Gold, M Tintora. 2013. Sex and gender issues in multiple sclerosis. Ther. Adv. Neurol. Disord. 6: 237–248.

10. J Sellner, J Kraus, A Awad, R Milo, B Hemmer, O Stüve. 2011. The increasing incidence and prevalence of female multiple sclerosis-A critical analysis of potential environmental factors. Autoimmun. Rev. 10: 495–502.

11. SR Murúa, MF Farez, FJ Quintana. 2022. The immune response in multiple sclerosis. Annu. Rev. Pathol. 17: 121–139.

12. JF Cryan, et al. 2019. The microbiota-gut-brain axis. Physiol. Rev. 99: 1877–2013.

13. A Amaan, G Prekshi, S Prachi. 2024. Microbiome-gut-brain axis: AI insights. Insights Biol. Med. 8: 001–010.

14. S Jangi, et al. 2016. Alterations of the human gut microbiome in multiple sclerosis. Nat. Commun. 7: 12015.

15. E Sefik, et al. 2015. Individual intestinal symbionts induce a distinct population of RORγ+ regulatory T cells. Science 349: 993–997.

16. A Ordoñez-Rodriguez, P Roman, L Rueda-Ruzafa, A Campos-Rios, D Cardona. 2023. Changes in gut microbiota and multiple sclerosis: a systematic review. Int. J. Environ. Res. Public Health 20: 4624.

17. Z Li, et al. 2022. Differences in alpha diversity of gut microbiota in neurological diseases. Front. Neurosci. 16: 879318.

18. BD Wagner, et al. 2018. On the use of diversity measures in longitudinal sequencing studies of microbial communities. Front. Microbiol. 9: 1037.

19. RV Alvarez, NM Vidal, GA Garzon-Martínez, LS Barrero, D Landsman, L Mariño-Ramírez. 2017. Workflow and web application for annotating NCBI BioProject transcriptome data. Database (Oxford) 2017: bax008.

20. E Bolyen, et al. 2019. Reproducible, interactive, scalable and extensible microbiome data science using QIIME 2. Nat. Biotechnol. 37: 852–857.

21. K Katoh, J Rozewicki, KD Yamada. 2019. MAFFT online service: multiple sequence alignment, interactive sequence choice and visualization. Brief. Bioinform. 20: 1160–1166.

22. MN Price, PS Dehal, AP Arkin. 2009. FastTree: computing large minimum evolution trees with profiles instead of a distance matrix. Mol. Biol. Evol. 26: 1641–1650.

23. AD Willis. 2019. Rarefaction, alpha diversity, and statistics. Front. Microbiol. 10: 2407.

24. C Ricotta, et al. 2019. Rarefaction of beta diversity. Ecol. Indic. 107: 105606.

25. C Lozupone, R Knight. 2005. UniFrac: a new phylogenetic method for comparing microbial communities. Appl. Environ. Microbiol. 71: 8228–8235.

26. RW Johnson. 2022. Alternate forms of the one-way ANOVA F and Kruskal-Wallis test statistics. J. Stat. Data Sci. Educ. 30: 82–85.

27. MJ Anderson, DCI Walsh. 2013. PERMANOVA, ANOSIM, and the Mantel test in the face of heterogeneous dispersions: what null hypothesis are you testing? Ecol. Monogr. 83: 557–574.

28. NA Bokulich, et al. 2018. Optimizing taxonomic classification of marker-gene amplicon sequences with QIIME 2’s q2-feature-classifier plugin. Microbiome 6: 17.

29. S Weiss, et al. 2017. Normalization and microbial differential abundance strategies depend upon data characteristics. Microbiome 5: 27.

30. SN Choileáin, M Kleinewietfeld, K Raddassi, DA Hafler, WE Ruff, EE Longbrake. 2020. CXCR3+ T cells in multiple sclerosis correlate with reduced diversity of the gut microbiome. J. Transl. Autoimmun. 3: 100032.

31. NS Elsayed, et al. 2023. Genetic risk score in multiple sclerosis is associated with unique gut microbiome. Sci. Rep. 13: 16269.

32. M A, et al. 2023. Mediterranean diet and associations with the gut microbiota and pediatric-onset multiple sclerosis: a trivariate analysis. Res. Square (preprint): RS-2540052.

33. MM Mashraqi, et al. 2023. Molecular mimicry mapping in Streptococcus pneumoniae: cues for autoimmune disorders and implications for immune defense activation. Pathogens 12: 857.

34. RA Flaherty, JM Puricelli, DL Higashi, CJ Park, SW Lee. 2015. Streptolysin S promotes programmed cell death and enhances inflammatory signaling in epithelial keratinocytes during group A Streptococcus infection. Infect. Immun. 83: 4118–4133.

35. IC Chikanza, S Trollip, LI Sakkas. 2024. Regulatory T cells and autoimmunity. In: Regulatory T Cells and Autoimmune Diseases. pp. 41–56.

36. R Abdugheni, et al. 2022. Metabolite profiling of human-originated Lachnospiraceae at the strain level. iMeta 1: e58.

37. M Anshory, et al. 2023. Butyrate properties in immune-related diseases: friend or foe? Fermentation 9: 205.

38. TL Montgomery, et al. 2024. Identification of commensal gut microbiota signatures as predictors of clinical severity and disease progression in multiple sclerosis. Sci. Rep. 14: 10211.

39. AM Mincic, M Antal, L Filip, D Miere. 2024. Modulation of gut microbiome in the treatment of neurodegenerative diseases: a systematic review. Clin. Nutr. 43: 1832–1849.

